# SureTypeSC - A Random Forest and Gaussian Mixture predictor of high confidence genotypes in single cell data

**DOI:** 10.1101/393256

**Authors:** Ivan Vogel, Robert C. Blanshard, Eva R. Hoffmann

## Abstract

**Motivation:** Accurate genotyping of DNA from a single cell is required for applications such as *de novo* mutation detection, linkage analysis and lineage tracing. However, achieving high precision genotyping in the single cell environment is challenging due to the errors caused by whole genome amplification. Two factors make genotyping from single cells using single nucleotide polymorphism (SNP) arrays challenging. The lack of a comprehensive single cell dataset with a reference genotype and the absence of genotyping tools specifically designed to detect noise from the whole genome amplification step. Algorithms designed for bulk DNA genotyping cause significant data loss when used for single cell applications.

**Results:** In this study, we have created a resource of 28.7 million SNPs, typed at high confidence from whole genome amplified DNA from single cells using the Illumina SNP bead array technology. The resource is generated from 104 single cells from two cell lines that are available from the Coriell repository. We used mother-father-proband (trio) information from multiple technical replicates of bulk DNA to establish a high quality reference genotype for the two cell lines on the SNP array. This enabled us to develop SureTypeSC - a two-stage machine learning algorithm that filters a substantial part of the noise, thereby retaining the majority of the high quality SNPs. SureTypeSC also provides a simple statistical output to show the confidence of a particular single cell genotype using Bayesian statistics.

## 1 Introduction

Single cell genomics is an umbrella term for genotyping of individual cells from a heterogeneous population. The deconvolution of mixed populations allows detection of genetic diversity within a population of cells. Applications cover many disciplines from sequencing the complete genomes of microorganisms that are challenging to culture in the laboratory to *de novo* mutation detection in tumour cells (Huang et al., 2015). Detecting genomic changes in single cells is a sensitive procedure, complicated by the often rare, unique and precious nature of the starting material, such as during genetic testing of human embryos for diagnostic purposes.

Unlike sequencing of bulk DNA, single cell sequencing requires a whole genome amplification (WGA) step to generate sufficient material for genotyping by next-generation sequencing (NGS) or single-nucleotide polymorphism (SNP) array (Gawad et al., 2016). A typical human cell contains 8-16 pg nuclear DNA that must be amplified to meet the input requirements for PCR-free sequencing (1 μg) or SNP array analysis (400 ng). The efficacy of genotyping from a single cell is critically dependent on the WGA method. Genome coverage, replication fidelity and the level of technical noise, such as systematic or stochastic amplification bias, are the main features considered when choosing the WGA method. However, all WGA methods deteriorate the signal from single cells. The signal deterioration potentially carries two risks: (a) sub-optimally amplified signal can lead to a complete loss of information about a particular locus, and (b) uneven signal amplification of two alleles at a heterozygous locus can result in an erroneous homozygous genotype call. The latter is called allele drop out (ADO) and affects up to 30% of SNPs from a single cell (Blanshard et al., 2018).

After WGA, the amplified DNA can either be sequenced or analysed by SNP arrays. There are several tools for genotype calling from WGA amplified DNA after sequencing (Zafar et al., 2016; Bohrson et al., 2017, bioRxiv; Lodato et al., 2015; Bae et al., 2018). In theory, sequencing can detect genomic changes from single base mutations to whole chromosome imbalances (Huang et al., 2015; Gawad et al., 2016). However, the sequencing depth required for applications such as detection of rare variants in a population of cells is limited by the number of cells that can be assessed, making sequencing impractical. In contrast, although SNP arrays are limited by assessing only a sample of the genome, the technology allows the analysis of a wide range of genetic variants with good coverage in a fast and cost-efficient manner. SNP arrays have been successfully applied to WGA DNA for discovery of new mutations, especially larger deletions that can cause loss of heterozygosity in cancers (Wong et al., 2004, Leung et al., 2002). They are also used in linkage analysis to screen preimplantation embryos for the presence or absence of monogenetic variants that underlie serious genetic disorders. This is referred to as preimplantation genetic diagnosis or preimplantation genetic testing for monogenic disorder, PGT-M. This makes it critical that genome coverage is high and that the SNPs are typed with high precision (Handyside et al. 2010; Natesan et al., 2014; Zamani et al., 2015). SNP arrays have also been used to infer the parental origin of chromosome imbalances in human preimplantation embryos as well as recombination and segregation patterns in meiosis after WGA (McCoy et al., 2015; Ottolini et al., 2015). The unique genomic arrangement of SNPs that occur naturally as haplotype blocks, or can be induced experimentally, also allow lineage tracing in a variety of organisms, including by SNP array analysis (Woodworth et al., 2017; Ottolini et al., 2017).

There is a plethora of tools and algorithms currently available for genotyping bulk DNA from SNP arrays (Ritchie et al., 2011; Li et al., 2012). These algorithms are optimized for SNPs on the array and perform very well in terms of both call rates and sensitivity. However, an algorithm that is specifically designed for single cell variant calling using SNP arrays is currently missing. This is important because it is unclear how well the genotyping platforms deal with the biases introduced by whole-genome amplification of DNA from single cells. One solution is to include only SNP calls that are similar in properties to those from bulk DNA. This, however, causes a substantive loss of data (Zamani et al., 2015). It is also unclear how accurate genotyping is after the whole-genome amplification.

Genotyping from SNP arrays relies on the detection of emission intensities (X and Y). Thus, when both X and Y are above a certain threshold, the genotype is inferred as heterozygous (AB). In contrast, when only X or Y is detected above a certain threshold, a homozygous genotype is assigned (AA or BB). Current genotyping algorithms are based on two distinct approaches. Model-based algorithms do not require a training data set and assume that every SNP can be modeled from a linear combination of multivariate components (Teo et al., 2007, Giannoulatou et al., 2008). Reference-based algorithms perform genotyping based on a comprehensive database of reference variants. Parameters of these algorithms are inferred from a training dataset (*e.g.* the HapMap population, International HapMap Consortium, 2005) and are used for normalization of the raw data (Ritchie et al., 2009) or provide a confidence measure of the genotype (Kermani, 2008). The training of the parameters can be performed via supervised machine learning methods, in particular neural networks (Kermani, 2008).

Here, we present a comprehensive database of 104 single cell samples from two different cell lines that we SNP-typed and compared with their reference genotype. This allowed us to divide the data in two classes: (a) high quality single cell calls and (b) misclassified single cell calls caused by deteriorated signal. We used both classes to develop a two-layered algorithm that combines a supervised machine learning method with a model-based algorithm. We refer to this as SureTypeSC, which is able to identify the noise in the single cell data coming from erroneous whole genome amplification and then assign a probability score of a SNP being correctly genotyped.

## 2 Materials and Methods

### 2.1 Cell lines and molecular methods

We generated genotypes from whole genome amplified DNA (from single cells) or genomic DNA from bulk extraction using the Infinium Karyomapping Assay Kit (Illumina Inc., California, US). We obtained EBV-lymphoblastoid cell lines GM07228 and GM12878 from the NIGMS Human Genetic Cell Repository at the Coriell Institute for Medical Research, New Jersey, USA, and cultured these according to the supplier’s recommendation. All of the molecular methods and genotyping using GenCall for obtaining the SNP genotypes are provided in the Supplemental Methods.

### 2.2 MA transformation

The MA transformation is an application of the Bland-Altman transformation (Bland and Altman, 1999) that has been used extensively in the analyses of gene expression data when intensity values for two channels are compared using microarrays (red and green, referred to as X and Y, respectively).

Formally, we apply a linear-log transformation for every SNP, *i*, carrying a tuple of intensities (*x*_*i*_, *y*_*i*_) by calculating the values *m*_*i*_ and *a*_*i*_, as follows:

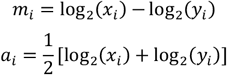

The M-feature has powerful discriminative ability to separate the three genotype clusters and is able to reduce variability between experiments and SNPs (Carvalho et al., 2007). The A-feature is a good general indicator of the signal quality (Ritchie et al., 2011).

### 2.3 Bioinformatics workflow for the machine learning algorithm

We developed a bioinformatics workflow with a supervised machine learning core that filters out the noise from the single cell data. The reference training intensities as well as validation intensities are first extracted from the intensity data (*.idat) files and subsequently genotyped using the GenCall algorithm implemented in GenomeStudio. The training data are then exported from GenomeStudio, transformed using the MA transformation and fitted to a two-layered machine learning model. The results are subsequently tested on a cross-validated data as well as on a set of independent single cell samples. The details of the workflow are shown in Fig. S1.

### 2.4 Training and validation datasets

We created a reference genotype for both single cell lines (GM07228 and GM12878) using parental information and multiple technical replicates from bulk DNA (Supplemental Methods). We subsequently compared our single cell datasets to the reference genotype. More specifically, for every candidate single cell call for SNP *i* and sample *s* we assigned a label: *l*_*i*,*s*_ ∈ {*True*, *False*}, depending on the match or mismatch with the corresponding reference genotype call. The training dataset is then a triplet (*m*_*i*,*s*_, *a*_*i*,*s*_, *l*_*i*,*s*_), where (*m*_*i*,*s*_, *a*_*i*,*s*_) are input features and *l*_*i*,*s*_ is the output feature. Note that we omit sample index *s* in further explanation, as we do not distinguish between the origins of SNPs in the training data set. We included all autosomal single cell calls with GenCall score above 0.01 (QC001) totaling 14,403,139 SNPs for training (GM07228) and 11,737,508 SNPs for validation (GM12878). Lowering the GenCall score threshold for accepting a SNP allowed as to include potentially poorly amplified SNPs and to capture the full error pattern. Table 1 and Table S2 give a detailed overview of the datasets used.

**Table 1.**
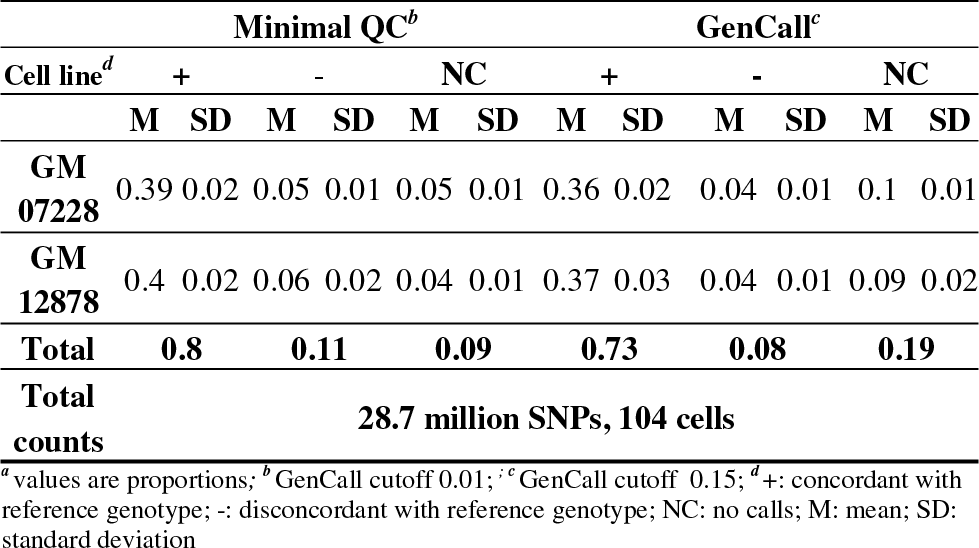
Summary of genotype calls from single cells^*a*^.

### 2.5 Supervised training using Random Forest

Random Forest is an ensemble supervised training method that is built from the collection (forest) of classification (decision) trees (Breiman, 2001). Each tree is trained on a different random subset of data and different subsets of input features. Although the training data only contain two input features (*M* and *A*), the preliminary analysis (Fig. 1B) suggests that the function that separates the erroneous clusters (red areas) from the correct calls (blue areas) is non-linear. Random Forest (RF) has the ability to fit different trees to different parts of the input space and therefore approximate a non-linear separating function resulting in increased classification accuracy. The class of miscalls (*l* = *False*) is usually a minority class and we therefore applied downsampling on the class of correct calls to tackle the class imbalance. We used the implementation of Random Forest from the scikit package (Pedregosa et al., 2011) for fitting the training data. We adjusted the following parameters of the algorithm:

- the number of trees was increased from 10 to 30; according to Oshiro et al., 2012, a theoretical upper limit is 128 trees and further increase in number of trees does not contribute to higher accuracy. However our data suggest that forests with more than 30 trees contribute minimally to the accuracy of the model but increase the size of the model substantially (data not shown);
- the number of features to consider when looking for the best split was set to two.

**Fig. 1.**
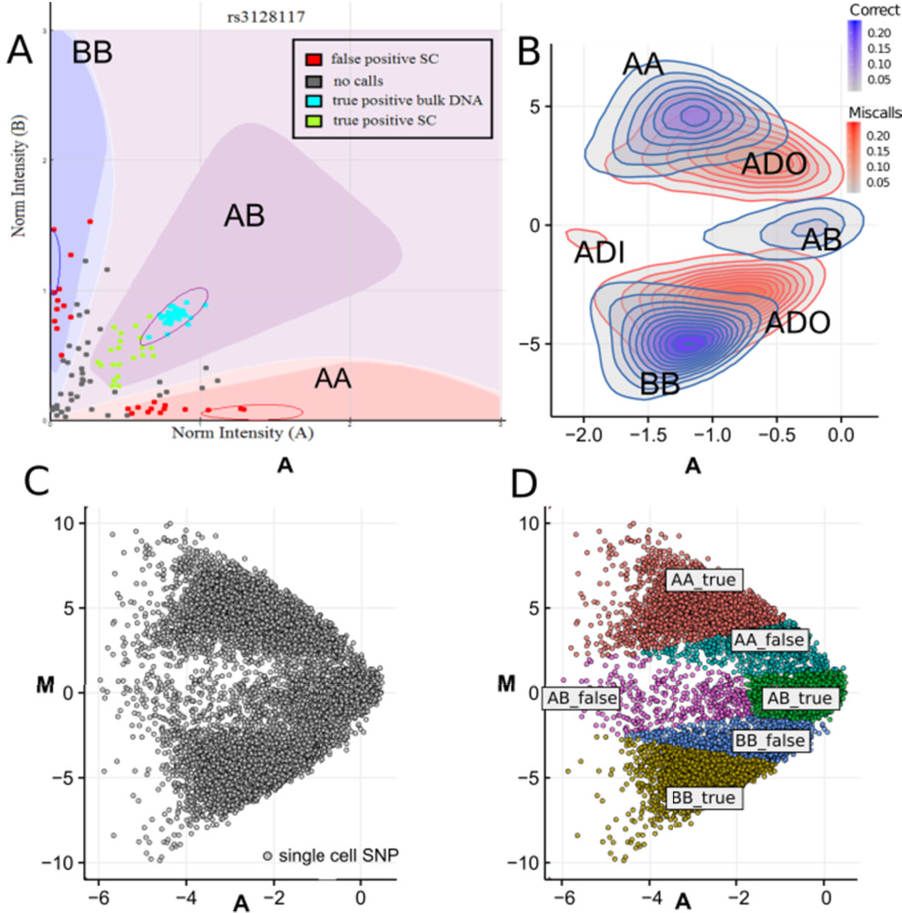
Signal-noise detection in whole-genome amplified DNA from single cells. (A) The GenCall algorithm in GenomeStudio classifies genotyping calls based on the normalized intensities of the X and Y channels (A and B allele, respectively). The genotyping space for homozygous AA calls is shown in red, heterozygous genotypes fall within the purple area and homozygous BB genotypes are in blue. The centroid of each genotyping space is shown as a circle. The genotyping space is specific to each SNP and based on bulk DNA. Cyan points: genotypes from bulk DNA, green points: correct genotypes from single cell, grey points: genotyping calls from single cells below the QC threshold of GenCall; red points: misclassified genotype from single cells. (B) Contour MA plot of all SNPs from one single-cell sample from GM07228; AA, BB and AB clusters are labeled accordingly. The probability density function was estimated using bivariate normal kernel from MASS library in R with default parameters. Each ring connects values with similar density levels and shades of blue/red show the likelihood of correct/incorrect genotype in that area (C) MA plot of 10,000 randomly selected SNPs from 10 single cell samples from GM07228. (D) Cluster labeling of randomly selected SNPs from (C) using SureTypeSC.

The prediction was evaluated in two ways - by stratified 10-fold cross-validation and with an independent single cell dataset. We used metrics that are commonly used in classifier evaluation as well as metrics that are specific for the single cell environment (Supplemental Methods)

### 2.6 Cluster correction using Gaussian Discriminant Analysis

The second stage of the algorithm is a Gaussian Discriminant Analysis (GDA) that formalizes the genotype clusters obtained from the RF step and potentially improves the precision and recall.

Let 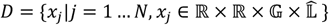 denote a set of N validation SNPs that were classified by the trained Random Forest, where 𝔾 = {AA, AB, BB}, 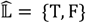. Therefore, 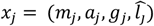 is a quadruplet of the logarithmic difference, logarithmic average, genotype predicted by GenCall (QC 0.01) and class prediction by RF at the *j*-th SNP. We assume that both the positive (T) and negative (F) classes, which are represented by pairs *d*_*i*_ = (*m*_*i*_, *a*_*i*_), come from mixtures of multivariate normal distributions. Based on this, we define the following system of Gaussian discriminants:

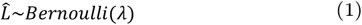

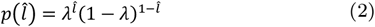

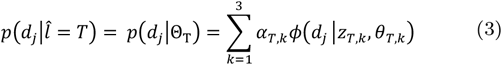

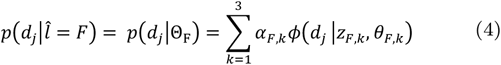

Where:

- *λ* denotes probability 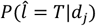
- *ϕ* is multivariate normal density function with parameters *θ*_*k*_ (with mean *μ*_*k*_ and covariance matrix Σ_k_)
- *z*_*k*_ is an indicator variable that denotes the genotype class, where 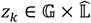
- *α*_*k*_ is the mixture component weight representing the probability that a random tuple (*m*_*j*_, *a*_*j*_) was generated by component *k*.

The complete set of parameters for the presented Gaussian discriminants is given as 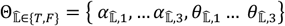

The log-likelihood function 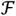 for classes from 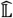 is defined as follows:

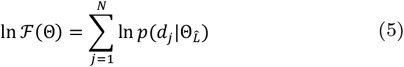

We use an Expectation Maximization algorithm (Dempster et al., 1977) to estimate the parameters 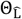 of the positive and negative class that maximize their log-likelihood function (Eq. 5). The EM algorithm is divided into an Expectation-Step (E-Step) and a Maximization-Step (M-Step). These are run in iterations separately for the positive and negative classes until convergence is reached. The detailed description of the algorithm is provided in Supplemental Methods.

After the parameters of both classes have been estimated by the EM algorithm, they are subjected to a second run. Here, the class membership 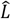 is hidden from the algorithm and every SNP is evaluated for both Gaussian discriminants using the following formula:

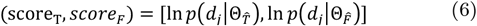

The final classification (membership to a positive or a negative class) is determined by higher value from the pair (score_T_, *score*_*F*_).

### 2.7 Scoring function

The key role of a genotyping algorithm is to report the likelihood of a certain genotype in form of a score or a posterior probability. Besides GenCall having its own scoring scheme, we used the following equations to estimate the probability of a certain SNP being correctly genotyped:

1. Random Forest: the score of a genotype of the *i*th SNP is given as a proportion of the trees in the forest that voted for a particular genotype being correct:

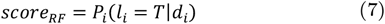
2. The scoring strategy of SureTypeSC is inferred from its second layer (GDA) as the class-conditional posterior probability of a genotype falling into positive class T:

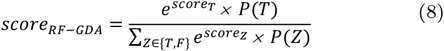

## 3 Results

### 3.1 Generation of 28.7 million high confidence SNPs from single cells

We typed nearly 28.7 million SNPs from 104 cells from two individuals (GM12878 and GM07228, Table 1 and Table S2) using the HumanKaryoMap-12 array (Illumina Inc., California, USA). To amplify the DNA from the single cells, we used multiple displacement amplification (MDA), a commonly used first-generation WGA method that relies on Phi (Φ) 29 polymerase. Its 3’→5’ activity allows proofreading and therefore improves the fidelity of amplification. This allows high precision genotyping with a mutation rate of 10^−7^ − 10^−9^. Furthermore, the ability to displace secondary DNA structures, such as hairpin loops that would cause other polymerases to stall or dissociate from the template DNA, allows the amplification of long DNA fragments (2-10 kb) (Blanshard et al., 2018; Dean et al., 2002).

### 3.2 Noise characterization of genotypes from single cells

To characterize the noise associated with genotyping from whole-genome amplified DNA from single cells, we compared the 28.7 million SNP genotypes from the two single cell datasets to their reference genotypes obtained from bulk, genomic DNA. To this end, we created high confidence reference genotypes from bulk DNA using nine independent bulk DNA samples hybridized against the HumanKaryomap-12 array and inferred genotypes using either the full parental information (GM07228, Supplemental Methods) or multiple technical replicates of bulk DNA and sequence data (GM12878, Supplemental Methods and Eberle et al., 2017). This allowed us to identify 264,269 SNPs for GM07228 and 270,681 for GM12878 (95.8% and 97.9 % of autosomal SNPs, respectively) on the HumanKaryomap-12 array that called correctly in every replicate from bulk DNA. From these, we generated high confidence reference genotypes.

Using the standard QC cutoff from GenCall (0.15), 73% SNPs (20.9 million) from the two single cell datasets called correctly, whereas 8% SNPs (2.36 million) were not concordant with the reference genotype (Table 1, Table S2). 19% SNPs gave ‘no calls’ (5.05 million; Table 1, Table S2), having failed to fall within the genotype clusters defined by bulk, DNA genotypes (Fig. 1A). The true positive rate was higher when we used a minimal QC (0.01) compared to the standard QC of GenCall (39%, SD=0.02% and 36%, SD=0.02%, respectively, for cell line GM07228 and 40%, SD=0.02% to 37%, SD=0.03% for GM12878, Table 1). These differences in true positive rates are statistically significant (p<0.0001, Fig. S2). In total for both datasets, the GenCall algorithm rejects about 7% of correctly genotyped SNPs from WGA DNA and increases precision by 2% (Table S2). We also listed call rates and error rates of individual cells and chromosomes from GM12878 and GM07228 (Table S3).

We displayed the pattern of the noise from the genotyping of SNPs from WGA DNA from single cells by first transforming the fluorescence intensities (X and Y) of each SNP into the logarithmic difference *M* and logarithmic average *A* (MA plot; Fig. 1B, C, D). At this stage of the workflow (“Building training dataset”, Fig. S1) we were able to observe the error pattern in the single cell data and display it in the form of contour plots (Fig. 1B). Three clusters of miscalls (false positives) became apparent in the single cell data. Two clusters were from allele drop out (ADO), where AB genotypes were incorrectly genotyped as AA or BB. A smaller cluster of allele drop in (ADI) also appeared. The ADI cluster was clearly separated from the true AB genotypes. Most of the errors, however, occur in the transition area between AB to AA or AB to BB (ADO) but nevertheless suggest good separability of the correct calls from miscalls, since the centers of the clusters are non-overlapping (Fig. 1B).

### 3.3 Design and implementation of the SureTypeSC algorithm

The characterization of the patterns of noise in a comprehensive dataset allowed us to employ a supervised machine learning method to classify and separate high quality genotypes from miscalls in the WGA DNA from single cells (Fig. S1). We combined a non-parametric (Random Forest) and parametric method (Gaussian Discriminant Analysis) and developed a scoring strategy that assigns probabilities that a specific SNP from a single cell dataset has been correctly genotyped (Methods, Eq. 7 and 8). Using a Random Forest prevents over-fitting of the data and provides good estimates of the positive and negative classes for the Gaussian discriminant analysis (Methods). We implemented the RF-GDA and the testing procedures in Python using the scikit library (Pedregoza et al., 2014) and pandas (McKinney, 2010). An example of a division of the feature space consisting of *M* and *A* by the RF-GDA algorithm is shown in Fig. 1D. Collectively, we refer the single layers (that can be implemented on their own) and the combined layers (RF-GDA) as SureTypeSC. The output from SureTypeSC is compatible with GenomeStudio and allows the user to import the results of the analysis back to GenomeStudio for further investigation.

### 3.4 Cross-validation of SureTypeSC

To assess whether our algorithm captures noise from the WGA and to exclude the possibility of overtraining, we first ran stratified 10-fold cross-validation on the single cell dataset from cell line GM07228. The dataset is imbalanced and mistyped SNPs are the minority class. We therefore used stratification to ensure that every fold contains both correctly genotyped and mistyped SNPs. In every iteration, we trained the algorithm on nine folds (27,445-27,772 SNPs) and used the 10^th^ fold for testing. To tackle the imbalance problem, we always balanced the training fold by downsampling the correctly genotyped SNPs. We evaluated the performance of every testing fold with the single layers individually (RF or GDA) as well as with a combination of them (RF-GDA). We scored the genotypes of all algorithms using the GenCall score or equations 7 or 8 (Methods). Consistent with random sampling of the SNPs, the mean performances of all algorithms have narrow confidence intervals (at 95%), which suggests that the algorithms are invariant to SNP selection (Table S4). Pairwise comparison of the algorithms using paired t-test shows that both, GDA and RF-GDA outperform GenCall in precision at similar recall (mean difference 4%, p < 0.0001, Table S5). Note that the RF outperforms GenCall in precision as well (mean difference 4.7%, p < 0.0001, Table S5), but has a lower performance in other metrics (Table S4). RF-GDA is more accurate than GenCall (mean difference 3.5%, p < 0.0001) and has a higher f1-score (mean difference 2%, p < 0.0001, Table S5).

### 3.5 Validation of SureTypeSC on an independent dataset

We next addressed how well our algorithms and GenCall performed on an independent dataset. To this end, we used the SNP genotypes obtained from 58 single cells after WGA from cell line GM07228 for training and the SNP genotypes obtained from WGA DNA from 46 single cells from a different cell line, GM12878 (Table 1, Table S2), for testing (‘tester set’). The genotyping data from the tester set were obtained at an independent time, with different batches of WGA reactions and genotyping arrays. This avoids systematic errors introduced by the chemistry used to obtain the genotypes. The training and validation scenarios are summarized in Fig. S3.

We first evaluated the performance of GenCall and SureTypeSC (RF, GDA and the RF-GDA) separately for heterozygous and homozygous regions using ROC and Precision-Recall curves (Fig. 2, Fig. S4). These metrics gave us visual insight into overall performance of the classifiers, invariant to the score cutoffs used. For the heterozygous calls, the RF-GDA outperforms all tested algorithms, which is also quantified by the ROC-AUC score (Table 2). Whereas GenCall achieves a 74% ROC-AUC score on average, this is increased to 86%, 87% and 92% for RF, GDA and RF-GDA, respectively (p < 0.0001, Table S6). For the homozygous regions, the RF outperforms GenCall at all points of the ROC and Precision-Recall curves, which is supported by the increase in the ROC-AUC score from an average of 67% (GenCall) to 81% for the RF (Table 2). This is further increased with the GDA or RF-GDA (both 83%). Interestingly, at a precision of 93%, the RF curve crosses that of the GDA and RF-GDA and recalls more true positive homozygous calls (Fig. S4A). This suggests that the RF alone might be a good option if higher recall is required at the costs of lower precision, which is nevertheless higher than GenCall in the homozygous regions. GenCall crosses the Precision-Recall curve of the RF-GDA at a precision around 88% and recalls more true positives (Fig. S4A). This is, however, very close to a recall of 100%, which also means accepting all calls without any significant filtration.

**Table 2.** Performance of the genotyping algorithms on independent dataset 12878^a^.

| Alg.<br>Metrics | GenCall<sup>b</sup> |  | RF |  | GDA |  | RF-GDA<sup>g</sup> |  |
| --- | --- | --- | --- | --- | --- | --- | --- | --- |
|  | het | homo | het<sup>c</sup> | homo<sup>d</sup> | het<sup>e</sup> | homo<sup>f</sup> | het | homo |
| accuracy | 0.68 ± 0.01 | 0.86 ± 0.012 | 0.71 ± 0.013 | 0.88 ± 0.008 | 0.63 ± 0.009 | 0.85 ± 0.01 | 0.84 ± 0.014 | 0.85 ± 0.01 |
| f1-score | 0.8 ± 0.01 | 0.92 ± 0.007 | 0.82 ± 0.012 | 0.93 ± 0.005 | 0.76 ± 0.011 | 0.91 ± 0.007 | 0.91 ± 0.011 | 0.91 ± 0.007 |
| precision | 0.97 ± 0.01 | 0.89 ± 0.009 | 0.99 ± 0.001 | 0.91 ± 0.008 | 0.99 ± 0.001 | 0.92 ± 0.008 | 0.99 ± 0.001 | 0.92 ± 0.008 |
| recall | 0.68 ± 0.01 | 0.96 ± 0.005 | 0.7 ± 0.017 | 0.96 ± 0.001 | 0.61 ± 0.013 | 0.9 ± 0.005 | 0.84 ± 0.017 | 0.9 ± 0.006 |
| ROC-AUC score | 0.74 ± 0.01 | 0.67 ± 0.015 | 0.86 ± 0.004 | 0.81 ± 0.012 | 0.87 ± 0.005 | 0.83 ± 0.013 | 0.92 ± 0.004 | 0.83 ± 0.012 |

<sup>a</sup> values are mean proportions over 46 cells +/- confidence interval at 95%; <sup>b</sup> GenCall score threshold 0.15; <sup>c</sup> Random Forest score threshold 0.6 and <sup>d</sup> 0.15; <sup>e</sup> Gaussian Discriminant Analysis score threshold 0.8 and <sup>f</sup> 0.5; <sup>g</sup> RF-GDA score threshold 0.15

**Fig. 2.**
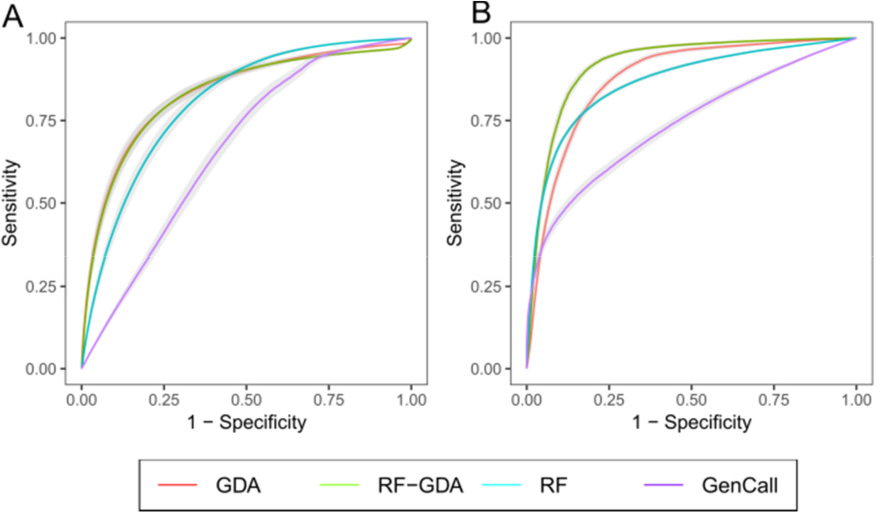
SureTypeSC improves the performance for single cell genotyping. ROC curve for homozygous (A) and heterozygous SNPs (B) on GM12878. The points of the curves were created by applying different cutoffs of the classification scores. Data below a certain cutoff were classified as negative, otherwise positive.The bands along the curves represent 95% confidence intervals from the 46 single cells.

Next, we were interested in how our methods perform compared to GenCall with standard settings (QC 0.15). Standard GenCall recalls 68% of the true positive heterozygous genotypes with a precision of 97%. The RF-GDA has 84% recall, achieves 99% precision on average, and thus outperforms the standard GenCall in both precision and recall. At similar precision, the RF and GDA on their own recall fewer true positive heterozygous genotypes (Table 2). As expected, high precision and recall is reflected in a high harmonic mean of precision and recall (f1-score) for the RF-GDA (Table 2) and a high rate of correctly classified SNPs (accuracy, Table 2). GenCall recalls 96% of the true positive homozygous genotypes on average at precision 89%. At similar recall, the RF alone increases precision by 2.5% (p < 0.0001; Table S6). GDA and RF-GDA further improve precision, but at the cost of recall. Both methods achieve an average precision of 92% at 90% recall for the homozygous calls (Table 2). Recalling fewer true positives at higher precision causes a drop in the f1-score for GDA and RF-GDA. This is because recall declines much quicker than the precision increases (Fig. S4A). The effect of lower recall from the GDA and RF-GDA is also mirrored in the lower accuracy. As GDA and RF-GDA have higher precision, they are also more likely to reject correct SNPs, thereby decreasing the number of true positives.

The two-layered architecture, RF-GDA, generally outperforms its constituent single layers (RF or GDA alone). Combining the RF and GDA together is particularly advantageous in the heterozygous regions, where the RF-GDA performs better in all metrics (Fig. S5). This is due to sensitivity of the EM algorithm to outliers, which are effectively reduced in the RF step (Supplemental Material, Section 1.8). For the homozygous calls, the RF-GDA performs better than single RF and GDA in precision (mean difference 1.7% and 0.2% for RF and GDA, respectively, p < 0.0001, Table S6).

However, the single GDA has better ROC-AUC score, which is 0.2% higher in the GDA than in RF-GDA (p < 0.001, Table S6). The ROC curve in Fig. 2A and Precision-Recall curve in Fig. S4A confirm that the difference is minor, since the RF-GDA and GDA largely overlap. Collectively, the benefits of the two layered RF-GDA compared to its single layers is the maximized precision and recall for the heterozygous calls. There is a further benefit in the maximized precision in the homozygous calls at the relatively modest loss of true positive calls.

### 3.6 Genotyping confidence in the single-cell environment

Our observations suggest that SureTypeSC can effectively improve precision of both homozygous and heterozygous SNPs (on average, 99% for heterozygous calls and 92% for homozygous calls, Table 2). Precision can be further improved at the cost of recall, particularly for homozygous SNPs, as Fig. S4 suggests. We therefore adjusted both SureTypeSC and GenCall for high precision, recalling ~47% of the true positive SNPs. To compare their performance, we developed a simple statistical toolkit that shows a detailed view of confidence in AA, BB or AB calls using a transition matrix of posterior probabilities (Table S9). The posterior probabilities show the probability that a certain genotype from the single cell application is genotyped correctly compared to the reference genotype. Table S9 shows that compared to GenCall, SureTypeSC achieves major improvements of 8% and 7% confidence of AA and BB, respectively, and an improvement of 0.3% in confidence of an AB genotype.

### 3.7 Allele-drop out and allele-drop in rates are reduced using SureTypeSC

Incorrect genotype calls arise predominantly from imbalances in the allele frequencies generated during the chemical reaction when the whole genome is amplified. The deviation from a 1:1 allele ratio of heterozygous SNPs can lead to allele drop out (ADO). Analogously, mistyping of a homozygous SNP results in allele drop in (ADI). We calculated the ADO and ADI rates for GenCall and SureTypeSC at high precision using the transition matrices from Table S9 (Table 3; performances of single layers RF and GDA are shown in Table S10). At a call rate of 42% for GenCall and 39% for SureTypeSC, GenCall is able to decrease ADI 7 times and SureTypeSC 12.5 times compared to minimal filtering (GenCall QC 0.01). The ADO rate is decreased 1.5 times by GenCall and 5.6 times by SureTypeSC (Table 3). Although SureTypeSC outperforms GenCall and minimizes the error incidence, the loss of data is inevitable (call rate 39%, Table 3).

**Table 3.** Allele drop-in and allele drop-out.

|  | Min. QC<sup>a</sup> | GenCall<sup>b</sup> | RF-GDA<sup>c</sup> |
| --- | --- | --- | --- |
| ADI | 0.05 ± 0.01 | 0.007 ± 0.003 | 0.004 ± 0.0005 |
| ADO | 0.14 ± 0.009 | 0.096 ± 0.01 | 0.025 ± 0.004 |
| Call rate | 0.92 ± 0.003 | 0.42 ± 0.01 | 0.39 ± 0.01 |

<sup>a</sup>GenCall QC threshold 0.01; <sup>b</sup>GenCall QC threshold 0.87; <sup>c</sup>RF-GDA score threshold 0.75. Proportions and confidence intervals at 95%. are shown.

### 3.8 Proof-of-concept of biological inference

To provide proof-of-concept that SureTypeSC would improve biological insight when used for high precision (RF-GDA), we assessed copy number variants (CNVs) in human oocytes (Fig. S7; Ottolini et al., 2015). The loss of a chromosome or chromosome segment results in one cell with only A and B calls (no heterozygous SNPs). The loss, however, is obscured by ADI when using the standard GenCall algorithm (Fig. S7). SureTypeSC removes the ADIs (erroneous AB), increasing the certainty of the inference (loss of hetSNPs).

We were interested in whether we could use SureTypeSC with high precision to reveal biological variability within the tested cell line GM12878. We assumed that all variants in the cell line that do not match the reference genome (‘erroneous variants’) are allele drop outs or allele drop ins (Fig. S8A). We were curious whether some of these variants, however, could be real and therefore used to detect heterogeneity within a cell population. We chose SureTypeSC as this documented the best overall performance in terms of precision (Table 2, Table 3, Table S9, Table S10) and compared it to GenCall. We performed hierarchical clustering (Supplemental Methods) on raw data with minimal filtering (QC 0.01), data processed by GenCall (GenCall QC 0.87), and RF-GDA at high precision (Table 3). The hierarchical clustering reveals there are potentially four subpopulations of cells in GM12878 cell line that are invariant to the type of filtration used (Fig. S8B, C, D). The bootstrap analysis (Supplemental Methods), however, reveals that only the RF-GDA consistently gives four stable subpopulations (Jaccard mean bootstrap value for a cluster > 0.75, Hennig 2007). The unstable clusters present in the trees from the minimal filter (QC 0.01) and ‘high precision’ genotyping using GenCall suggest non-reproducible noise being transferred to the bootstrapped replicates that is removed by SureTypeSC.

## Discussion

Whereas there are specialised tools for single cell genotyping from next-generation sequencing data (Zafar et al., 2016; Bohrson et al., 2017, bioRxiv; Lodato et al., 2015; Bae et al., 2018), no such algorithms exist for genotyping data from SNP arrays. Instead, genotyping WGA DNA has relied on increasing the threshold of the genotyping algorithms, which causes a substantial data loss (Zamani et al., 2015).

In this study, we have typed nearly 30 million SNPs from 104 single cells from two independent cell lines and developed an algorithm to distinguish signal from noise in whole-genome amplified DNA. SureTypeSC consists of two layers – a Random Forest (RF) and a Gaussian Discriminant Analysis (GDA) that work singly or in a cascade. The cascade approach is particularly beneficial for heterozygous SNPs, as it improves both precision and recall compared to the single layers and GenCall. We observed an increase in recall from 68% in GenCall at standard QC (0.15) to 84% in SureTypeSC. Resolving most of the heterozygous SNPs makes SureTypeSC highly relevant and applicable when heterozygosities are needed, such as tag SNPs during linkage analysis of transmission of monogenic diseases and aneuploidy detection (Handyside et al., 2010; Natesan et al.,2014; Zamani et al.,2015). At the same time, SureTypeSC improves precision for both homozygous and heterozygous SNPs by 3% and 2%, respectively (Table 2, Table S8). Having high precision makes it feasible to explore rare events across populations of cells. This includes assessing clonal expansion in tumour evolution, linage tracing, or detecting rare *de novo* mutations such as large genomic rearrangements in single cells that are averaged out and lost in bulk analyses (Cooper et al., 2015; Lu et al., 2012; Chen et al., 2017; Wong et al., 2004, Leung et al., 2002). High precision is also needed to obtain high resolution at haplotype breakpoints, which is particularly important in diagnostics (Handyside et al., 2010; Natesan et al., 2014; Zamani et al., 2015). Our proof-of-concepts show that SureTypeSC is likely to improve diagnostics as well as biological inferences.

As running the single layers of SureTypeSC could be potentially beneficial, such as high recall in homozygous regions at lower precision by the RF alone, SureTypeSC always operates in both modes (cascade and single layers), simultaneously, and scores the genotypes with RF, GDA, as well as RF-GDA.

Analysing a large number of single cells allows the decomposition of heterogeneous populations. Understanding how single cells in a population contribute towards a ‘mosaic’, mixed SNP call is particularly important as use of SNP arrays increases in cytogenetics. Furthermore, having a robust algorithm of genotyping from WGA DNA from single cells improves the certainty of genotype calling when only few cells are available. This is important in both basic biomedical research as well as clinical settings such as in preimplantation genetic testing. We show that SureTypeSC can be used for both (Fig. S7, Fig. S8).

We have implemented SureTypeSC in two modes in terms of performance. Using the standard mode where precision and recall are balanced, SureTypeSC was able to successfully identify a chromosomal loss in the single cell oocyte data, where GenCall failed to reject SNP calls from the chromosomal region (Section 3.8; Fig. S8). Using a high precision mode, SureTypeSC, but not GenCall, was able to stably detect four subpopulations in the reference GM12878 cell line. Thus, SureTypeSC most likely revealed true heterogeneity within the single cell population. This allows the use of SNP arrays in the single cell environment to explore fine differences between closely related cells. This was previously not possible due to the low resolution of the SNP array in combination with the noise coming from WGA.

Although genotyping from SNP arrays cover only a fraction of the genome compared to next-generation sequencing, the cost of *de novo* genome assembly is prohibitive even for bulk, genomic DNA when assessing a large number of cells or samples. The sequencing depth, or coverage, needed in one recent reference genome assembly for the detection of *de novo* mutations was nearly 50× (Besenbacher et al., 2015). For single-cell applications, the coverage to accurately identify new mutations from the noise and bias introduced by the whole-genome amplification step is in excess of this (Behjati et al., 2014). Thus, SureTypeSC allows a cost-effective approach to improve genotype accuracy using SNP arrays.

## Acknowledgements

The following cell lines/DNA samples were obtained from the NIGMS Human Genetic Cell Repository at the Coriell Institute for Medical Research: GM07228, GM07224, GM07225 and GM12878. We thank Alex Bladon from Illumina Inc. for critical discussion and members of Hoffmann lab for comments on the manuscript.

