## Supplemental Methods, Supplemental Figures and Supplemental Tables for "SureTypeSC - A Random Forest and Gaussian Mixture predictor of high confidence genotypes in single cell data": Supplemental_Material.docx

**1. SUPPLEMENTAL METHODS.**

### 1.1 Cell lines

All procedures with cell lines were performed in a Class II Microbiological Safety Cabinet (CIIMSC) unless stated otherwise. Genomic DNA was extracted from cell pellets containing 3.0×10^6^ cells, using the QIAamp DNA Mini Kit, including the Appendix B protocol for cultured cells (QIAGEN, Hilden, Germany), according to the manufacturers’ instructions. Extracted DNA quality was assessed by NanoDrop 1000 spectrophotometer (Thermo Fisher Scientific Inc., Massachusetts, USA), prior to storage at −80 °C until use. Single cell isolation from cell lines was performed 24-48 hours following passage. Sterile filtered 1×PBS and 1×PBS + 0.1% PVP (clinical grade polyvinylpyrrolidone; Origio, Måløv, Denmark) isolation buffers were prepared fresh for each isolation session from stock solutions. Sterile, thin-walled 0.2 ml PCR tubes were labeled and primed with 3 µl 1×PBS. 0.9×10^6^ cells were harvested in the logarithmic phase of growth, and dead cells were removed using the MACS magnetic, microbead-based Dead Cell Removal Kit with MACS MS column separation (Miltenyi Biotech, Bergisch Gladbach, Germany) according to the manufacturers’ instructions and eluted into 1 ml 1×PBS + 0.1% PVP by pipette mixing. Cell viability was assessed prior to- and post-dead cell removal by staining with 0.4% trypan blue and counting under an Olympus CKX31SF inverted microscope (Olympus, Shinjuku, Japan). Cell solutions with a viability of >98% were used for immediate single cell isolation and kept at 4 °C. Cells were isolated by manual micromanipulation in 1×PBS + 0.1% PVP using a STRIPPER Micropipette and STRIPPER tips with a 125 µm lumen, with an aspiration volume of 1 µl (Origio, Måløv, Denmark), under an M125 C stereo microscope (Leica, Wetzlar, Germany) in an AirClean 600 microscope cabinet (AirClean Systems Inc., North Carolina, USA). Briefly, single cells were placed into individual 6 µl isolation drops before gentle washing by repeated aspiration in fresh 6 µl wash drops, followed by immediate tubing. The final sample volume in each tube was 4 µl. Between cell samples, micropipette tips were washed thoroughly in 1×PBS by aspiration. During isolations, sample tubes were stored on a −20 °C tube rack. An isolation buffer only (negative control) and 3-cell positive control were also collected. Upon completion of isolation, cell samples were transferred to −20 °C for storage prior to whole-genome amplification within 48 hours. Alternate cell lines were isolated on separate days using fresh preparations of all reagents.

### 1.2 Whole genome multiple displacement amplification

Sample amplification by multiple displacement amplification (SureMDA, Illumina Inc., California, USA) was performed according to the Infinium Karyomapping Assay Protocol Guide (Part: 15052710 RevB; Illumina Inc., California, USA). A human male genomic DNA amplification control was prepared to give 50 pg DNA in 4 µl kit-supplied 1×PBS. A 4 µl ’no template control’ (NTC) was prepared from 1×PBS only. All sample pipetting steps were performed over −20 °C tube racks in a CIIMSC in a designated pre-PCR laboratory. Reagents were dispensed on the inner wall of each sample tube above the sample droplet, prior to centrifugation at 280 × *g* for 5 sec. Immediately before use, single cell isolates and isolation controls were removed from −20 °C storage and centrifuged at 300 × *g* for 3 min. MDA was performed according to the manufacturer’s instructions, using an Applied Biosystems Veriti Thermal Cycler (ThermoFisher Scientific Inc., Massachusetts, USA). Samples were amplified for 2 hours at 30 °C and stored at −20 °C prior to resolution by gel electrophoresis and quantitation of dsDNA. Samples were resolved by gel electrophoresis on a 0.8% agarose gel in 1×Tris-borate-EDTA (TBE) buffer with 1×GelRed™ stain (Biotium Inc., California, USA), for 50 min at 120 V over 30 cm. Duplex DNA was quantified using Quant-iT dsDNA Assay Kit, High Sensitivity (ThermoFisher Scientific Inc., Massachusetts, USA).

### 1.3 Illumina Infinium II assay and BeadArray

Single cell MDA products and genomic DNA extracted from cell lines were genotyped using the Infinium Karyomapping Assay Kit (Illumina Inc., California, USA), according to the manufacturer’s instructions. Briefly, genomic DNA samples were adjusted to 50 ng/µl in 10 mM Tris-HCl, pH 8.5. Single cell MDA products were not diluted. Prepared DNA was hybridized to HumanKaryomap-12 v1.0 BeadChip Arrays at 48 °C for 16-20 hours. BeadChip arrays were washed using kit-supplied reagents before assembly into Te-Flow flow-through chambers, and inserted into a Tecan Chamber Rack (Tecan, Männedorf, Switzerland). Single base extension of Infinium oligonucleotide probes was performed at 44 °C, prior to denaturation with 95% formamide + 1 mM EDTA to remove the template DNA. Staining of biotin and dinitrophenyl (DNP) labelled nucleotides for signal amplification with two-colour master mix and anti-stain was performed at 32 °C. BeadChip arrays were immediately washed and coated before vacuum drying at >0.9 bar for 90 minutes. Processed arrays were scanned using either the iScan or NextSeq550 System (Illumina Inc., California, USA), and intensity data were analysed using GenomeStudio v2.0.2 software with Genotyping Module v1.9 (Illumina Inc., California, USA) for genotype calling.

### 1.4 Normalisation and Genotyping using GenCall

The GenCall algorithm is part of Illumina’s Genotyping module. First, raw intensity data from the two channels (X and Y) are normalized by removing outliers, adjusting for background noise and employing affine transformation on the data. The objective of the affine transformation is to translate and scale data towards the x and y axes for the two homozygous genotype clusters (AA and BB, respectively). The polar coordinates are fitted into predefined genotype clusters with their centroid predicted by a neural network. The neural network was trained on genomic bulk DNA from the Phase I HapMap Project (International HapMap Consortium, 2005) for three global populations with minor allele frequency (MAF) >0.05. These samples form a centroid as the default position to which the genotype is inferred by determining the nearest cluster. A confidence score that can range from 0 to 1 , known as the GenCall score (GC), is assigned to every SNP and is used as a measure of quality. The closer the genotype is to the center of the assigned cluster, the higher is the GenCall score. As a default, genotypes with GC values lower than 0.15 are considered false positive. The user has an option to generate new cluster files (’recluster’) using the user-derived data, however this step requires sufficient coverage of all three genotypes for every SNP – according to Illumina’s documentation this is approximately 100 individuals per locus per population (Illumina, 2014, available at https://www.illumina.com). Illumina’s algorithm and its default configuration are optimized for bulk, genomic DNA.

### 1.5 Generation of reference genotypes on the HumanKaryomap-12 v1.0 BeadChip Arrays

In order to create a reliable training data set of single cell SNPs, we first created a high quality reference genotype that would serve as ground truth for our observations. We operated with multiple replicates of bulk genomic DNA (nine replicates of maternal bulk DNA from GM07224, six replicates of paternal bulk DNA from GM07225 and nine replicates of proband bulk DNA that was also subjected to the single cell analysis from GM07228). To assess a reference genotype for the subject DNA, we created a consensus genotype for mother, father and proband by only accepting sites that show 100% concordance between the replicates. Furthermore we checked for correct parental inheritance patterns and excluded all proband’s genotypes that did not follow the rules shown in Table S1. This process excluded 4.15 % of the autosomal SNPs. For the testing dataset GM12878 we built a reference genotype by creating a consensus of five replicates of bulk DNA and only including SNPs that matched the genotyping from deep sequencing (Eberle et al., 2017), which excluded 2.1% of the autosomal SNPs.

### 1.6 Validation metrics

We used the following metrics for validation:

- Recall (or sensitivity): $\frac{TP}{TP+FN}$
- Precision: $\frac{TP}{TP+FP}$
- Specificity:$\frac{TN}{TN+FP}$
- F1 score as the harmonic mean of precision and recall
- Receiver Operating Characteristic curve (ROC), which shows sensitivity (true positive rate) as a function of 1-specificity (false positive rate). The points of the curve are created by calculating true positive rate and false positive rate for different threshold of the tested algorithm
- ROC-AUC score – area under the ROC
- Precision-Recall (PR) curve displays recall as function of precision. The points of the curve are created by calculating precision and recall curve at various thresholds of the tested algorithm.
- Posterior Probability Matrix is a custom-defined metric showing the posterior probabilities $P\left( g_{ref} | g_{sc} \right)$, where $g_{ref}$ is the reference call (in rows) and $g_{sc}$ is the single cell call (in columns); posterior probability in this context is a confidence measure of genotype$g_{sc}$ having a truth value of $g_{ref}$
- Allele drop-in (ADI) is an erroneous change from homozygous to heterozygous genotype and can by calculated from the Posterior Probability Matrix by applying: $P\left( AA | AB \right)+P(BB|AB)$
- Allele drop-out (ADO) is an erroneous change from heterozygous to homozygous genotype and can be calculated from the Posterior Probability Matrix by calculating $P\left( AB | AA \right)\times P\left( \mathrm{AA} | \hom\right)+P\left( \mathrm{AB} | \mathrm{BB} \right)\times P(BB|hom)$

TP, FN, TN and FP mean true positive, false negative, true negative and false positive respectively. The points of ROC and PR curve were drawn by applying various cut-offs of the algorithms.

### 1.7 Expectation Maximization algorithm

Expectation Maximization algorithm is part of the second layer of SureTypeSC (GDA). $N_{\hat{l}}$ is total number of SNPs in the particular class.

**Algorithm:**

For every SNP *i*	with label $\hat{l}$ in $\hat{\mathbb{L}}$:

1. E-step - calculate membership weights - probabilities of ($m_{i},a_{i}$) belonging to a cluster *k* using either initialization parameters $\Theta_{\hat{l}}^{init}$ if this is the first iteration, otherwise $\Theta_{\hat{l}}^{t+1}$

| $\omega_{i,k,\hat{l}}=p\left( z_{i,k,\hat{l}}=1 \vert d_{i},{\Theta_{\hat{l}}^{t}} \right)$=  $=\frac{\Phi\left( d_{i} \vert z_{i,k,\hat{l}}\theta_{k,\hat{l}} \right)\cdot\alpha_{k,\hat{l}}}{\sum_{m=1}^{K} \Phi\left( d_{i} \vert z_{i,m,\hat{l}},\theta_{m,\hat{l}} \right)\cdot\alpha_{m,\hat{l}}}$  for $1\leq k\leq3, 1\leq i\leq N.$ | (1) |
| --- | --- |

2. M-step
   - Calculate new component weights for the next iteration

$$\alpha_{k,\hat{l}}^{t+1}=\frac{\sum_{i=1}^{N_{\hat{l}}} \omega_{i,k,\hat{l}}}{N_{\hat{l}}}$$

(2)

- calculate new means for the next iteration

$$\mu_{k,\hat{l}}^{t+1}=\frac{\sum_{i=1}^{N_{\hat{l}}} \omega_{i,k,\hat{l}}d_{i}}{\sum_{i=1}^{N_{\hat{l}}} \omega_{i,k,\hat{l}}}$$

(3)

- calculate new covariances for the next iteration:

$$\Sigma_{k,\hat{l}}^{t+1}=\frac{\sum_{i=1}^{N_{\hat{l}}} w_{i,k,\hat{l}}\cdot(d_{i}-\mu_{k,\hat{l}}^{t+1}){(d_{i}-\mu_{k,\hat{l}}^{t+1})}^{T}}{N_{k,\hat{l}}}$$

(4)

at the end of the M-step we obtain new parameter estimates Θ^t+1^

3. Calculate log likelihood using Eq. 5 and if the relative change in the overall likelihood is smaller than a threshold, halt. Otherwise proceed with the E-step with parameters from Θ^t+1^.

Note that we use two different categories of weights - membership weights ($\omega$) and component weights ($\alpha$). Membership weights are related to a particular SNP, whereas component weights represent the relative call rate of a particular genotype.

### 1.8 Validation of the heterozygous components in the Gaussian mixtures

### The results show that the cascade structure of the RF-GDA is beneficial for the heterozygous SNPs and improves precision and recall compared to RF or GDA alone (Table 2, Fig. 2, Fig. S4). The difference in the implementation between GDA and RF-GDA is that GDA approximates the parameters from the ground truth data whereas RF-GDA approximates the parameters from the data labeled by RF (Fig. S3). To explain the differences in performance, we calculated the parameters of the heterozygous cluster from the ground truth (Table S7A) and extracted the parameters of the heterozygous component of the positive and negative class from the GDA model, when fitted on the ground truth data or on data predicted from RF or GDA (Table S7B). We also measured the data properties of the heterozygous cluster after prediction with RF or GDA (Table S7C), before the application of the second layer (EM algorithm from GDA). The results confirm that the heterozygous clusters in the ground truth dataset GM12878 and GM07228 are similar, as inferred from the similar eigen values and eigen vectors of the covariance matrices shown in Table S7A). In contrast, when fitting the GDA on the ground truth data (GM07228), the parameters of the positive and negative heterozygous components differ greatly from the ground truth (Table S7A and M1 and M2 in Table S7B). While covariance is close to zero in the ground truth data suggesting that the data mainly changes in the direction of the x axis (a-feature), it is much higher in the model M1 and M2. Furthermore, the high variance of the m-feature in the fitted model also indicates that the EM algorithm did not fit to the ground truth correctly. We demonstrate the problem of suboptimal fitting by plotting the ground truth distribution of the heterozygous cluster (Fig. S6A) and then distribution of the data generated from the model with parameters fitted on the ground truth (Fig. S6B). In contrast, when the GDA is fitted on the data predicted by RF, the parameters of the model generate a distribution that closely resembles the original ground truth data, particularly for the positive class (Fig. S6D, M3 and M4 in Table S7B). To explain this, we investigated the properties of the data derived after the RF prediction and prior to fitting with the EM algorithm (Models D1 and D2 in Table S7C). The histogram in Fig. S6C shows that the RF generates more negative labels and removes the typical tail of the original distribution for the positive class. Moreover, the RF also shifts the mean of the negative class and decreases the variance of the positive class, whichenables the EM algorithm implemented in GDA to generate an improved fit of the target distribution. As a final test, we measured the discriminative power of the fitted GDA models (Table S8) and found that the GDA fitted on data predicted by the RF has the highest outcome among all models (Supporting Classifier RF_7228 and RF_12878 in Table S8). Throughout the parameter testing, we always considered all combinations of the two ground truth datasets as well as using GDA in the first layer of the algorithm instead of RF. The presence of the tails in the distribution are likely due to truebiological variation in the individual cells from the cell lines. From the technical point of view, these are seen as outliers. Sensitivity to outliers (or generally tails in the distributions) is a known issue of the MLE algorithms. Our results show that processing the data through RF simplifies the distribution, thereby improving the fit of the EM algorithm of the heterozygous component of the positive and negative classes, resulting in increased discriminative power and thus performance for the heterozygous calls.

### 1.9 Exploring heterogeneity in GM12878

We performed high precise genotype filtering using GenCall and RF-GDA (SureTypeSC) of 46 cells from GM12878 (Methods). Both algorithms were adjusted for high recall (GenCall score 0.87, SureTypeSC score 0.75) To explore the heterogeneity in terms of concordance with the reference bulk DNA genome, we coded variant genotypes that corresponded the reference with one and variants not matching the reference with zero (similarly to building the training dataset, Methods). Similarly to Zafar et al., 2016, we imputed the missing genotypes with value 0.5 and used Hamming distance to calculate the pairwise dissimilarity between the 46 cells in GM12878. We then performed hierarchical clustering using hclust in R on all SNPs that contained at least one non-reference variant across the 46 cells. We included raw,non-filtered) genotypes (QC001) ,genotypes from GenCall (0.87), and SureTypeSC (0.75) and used Ward distance for hierarchical clustering. We assessed stability of the clusters by performing bootstrap analysis implemented in the fpc package in R. We subsequently labelled the clusters with clusterwise Jaccard bootstrap mean (calculated from 1000 replicates) indicating the stability of the cluster (Figure S8).

### 1.10 Training on balanced and unbalanced datasets

### We always balanced the training dataset prior to the training in both layers of SureTypeSC by downsampling the positive class. We tested the performance of the algorithms when using original imbalanced data and compared it to the performance of the balanced solution. As shown in Fig. S9C, balancing the dataset is particularly beneficial for the RF layer as it improves precision for homozygous genotype calls.

**2. SUPPLEMENTAL FIGURES.**

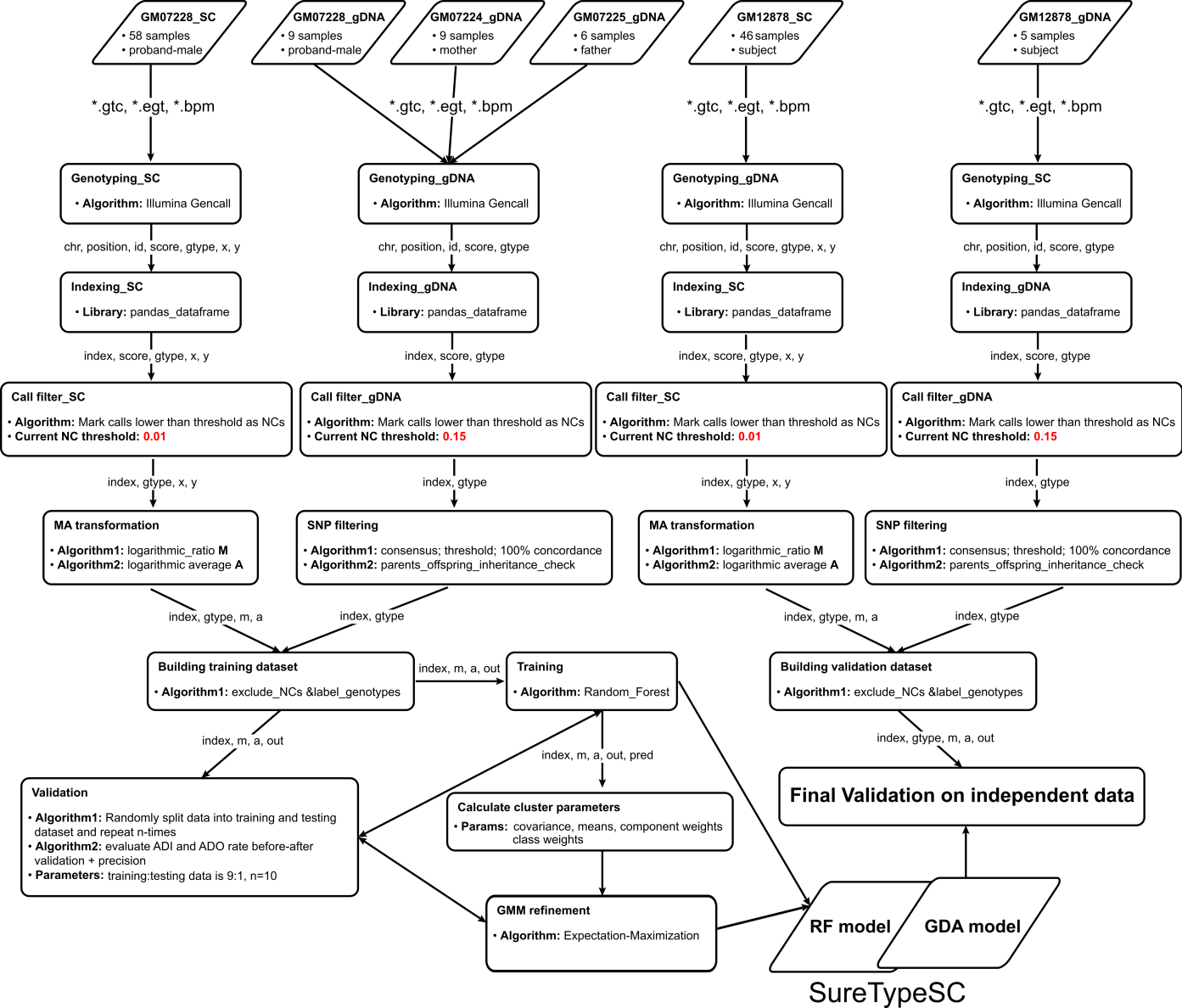

**Figure S1. Detailed workflow for the development of SureTypeSC.** 58 single cells from GM07228 were genotyped. High confidence genotypes were generated from the bulk DNA from the trio (paternal, maternal and son- GM07228). The parallel branch for testing 46 single cell genotypes from cell line GM12878 or as five bulk DNA was also set up. The intensities were stored in the Genotype Call Files (*.gtc), the information about SNPs and probe content is stored in BeadPool Manifest file (*.bpm). The cluster files (*.egt) carry the reference information for each locus. All files were available prior to the single-cell genotyping analysis (the intensity files are the final product of the BeadChip scanning procedure and *.egt and *.bpm are available online from Illumina). The data is was exported from GenomeStudio, indexed using chromosomal position (chromosome and position) and SNP ID (id) in a dataframe structure from Pandas library. The MA-transformed data are were fitted to the two-layered machine learning model and validated using both independent and cross-validated datasets.

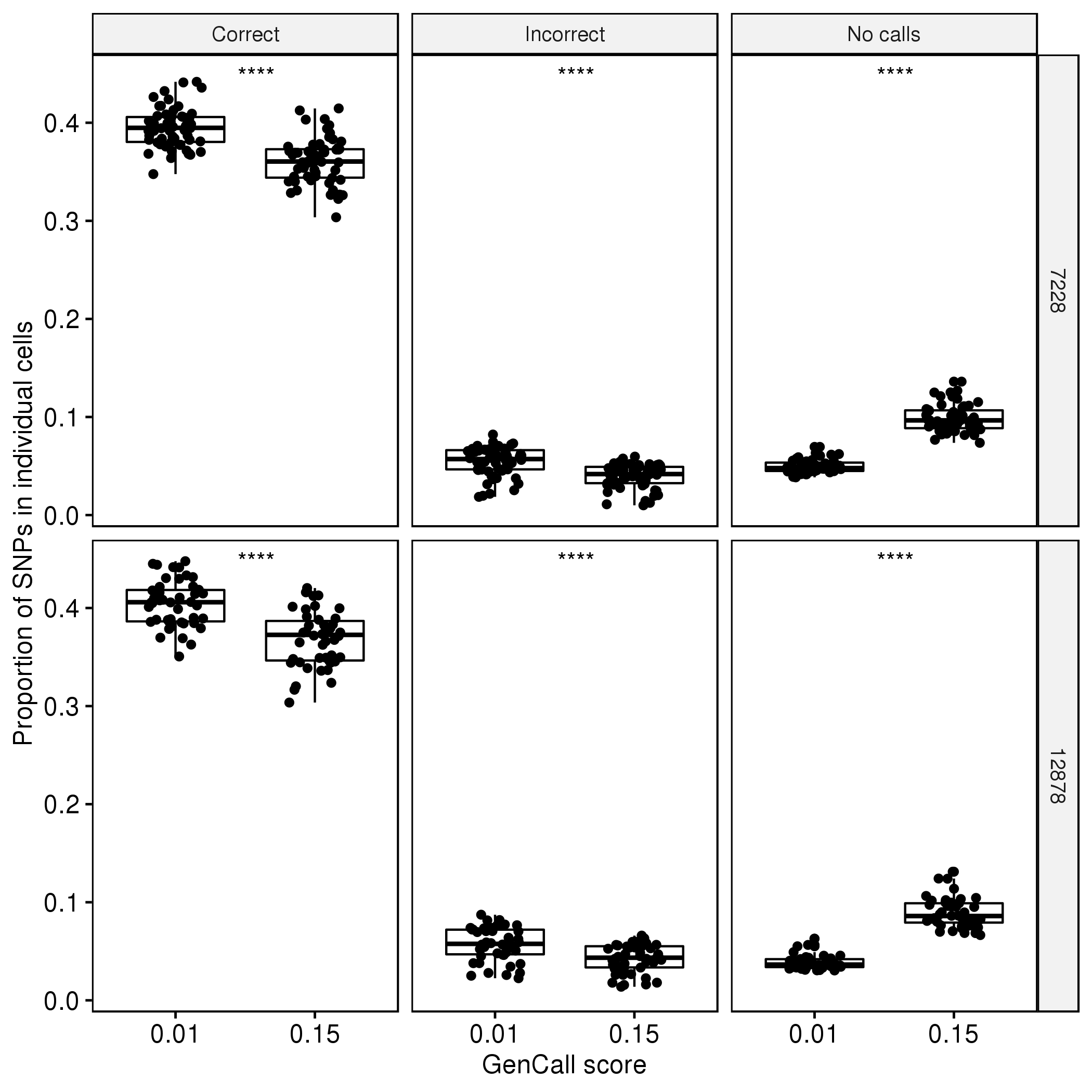

**Figure S2. Rates of correct calls, incorrect calls and no-calls for 58 individual cells of GM7228 and 46 cells of GM12878.** Each dot represents one cell. The proportions of SNPs within each single cell, that were correctly typed, incorrectly typed, or not typed (rejected due low GenCall score) are shown in the plots from left to right. Paired t-test was used to draw differences between cells typed with GenCall score 0.01 and GenCall score 0.15.

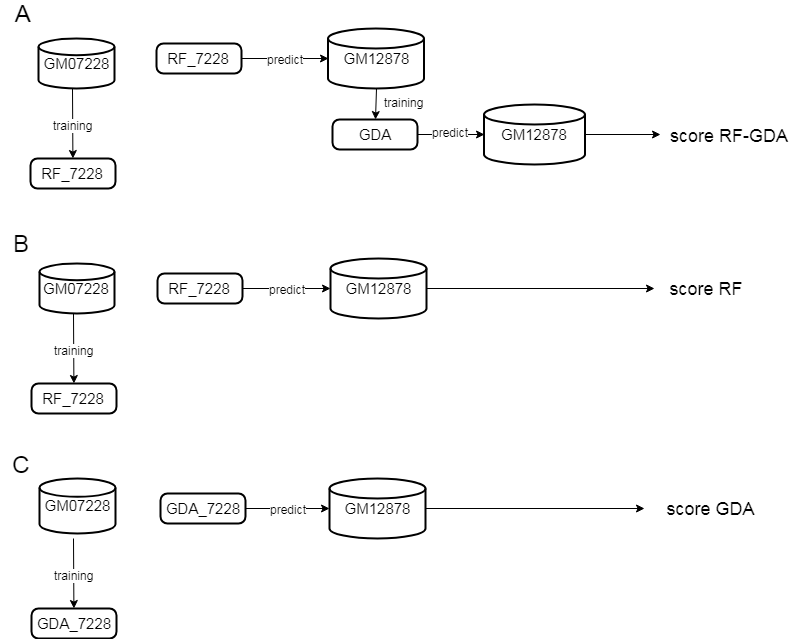

**Figure S3. Training and testing strategy for SuretypeSC.** (A) The RF was trained on ground truth data from GM07228 and used to predict the values of GM12878. The GDA was used fit the predicted values of GM12878 (B) The RF was trained on ground truth data from GM07228 and used for prediction and scoring on the testing data, GM12878. (C) The GDA trained on ground truth from GM07228 and prediction and scoring took place on the testing data, GM12878.

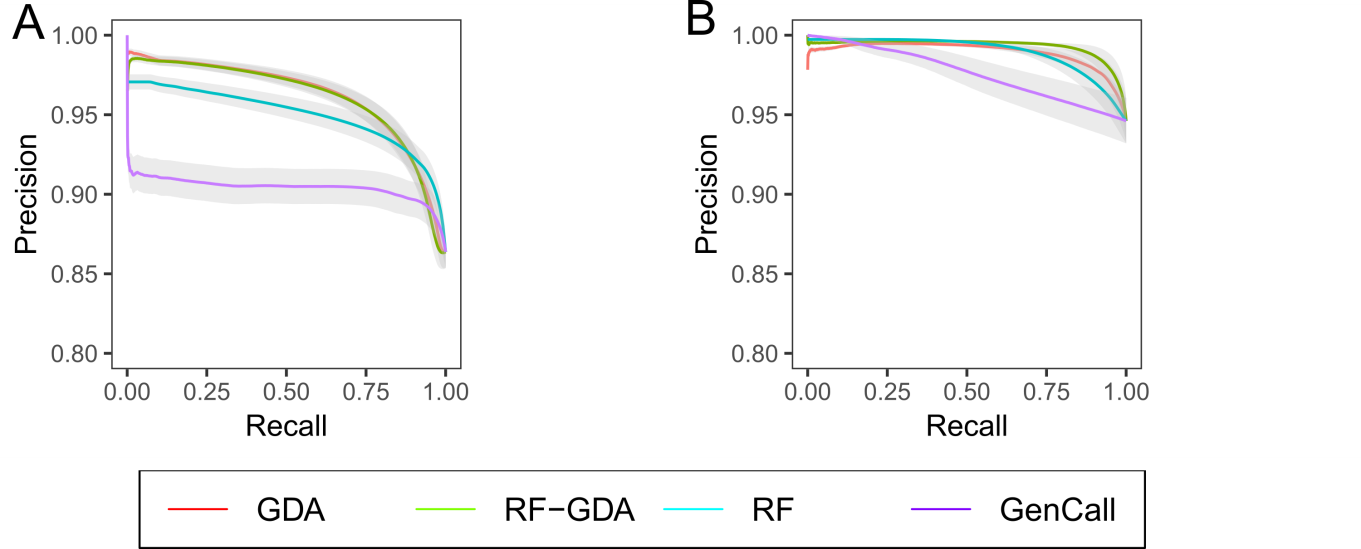

**Figure S4 Precision-Recall curves for homozygous (A) and heterozygous SNPs (B).** The points were drawn by applying different cut-offs to the algorithms.

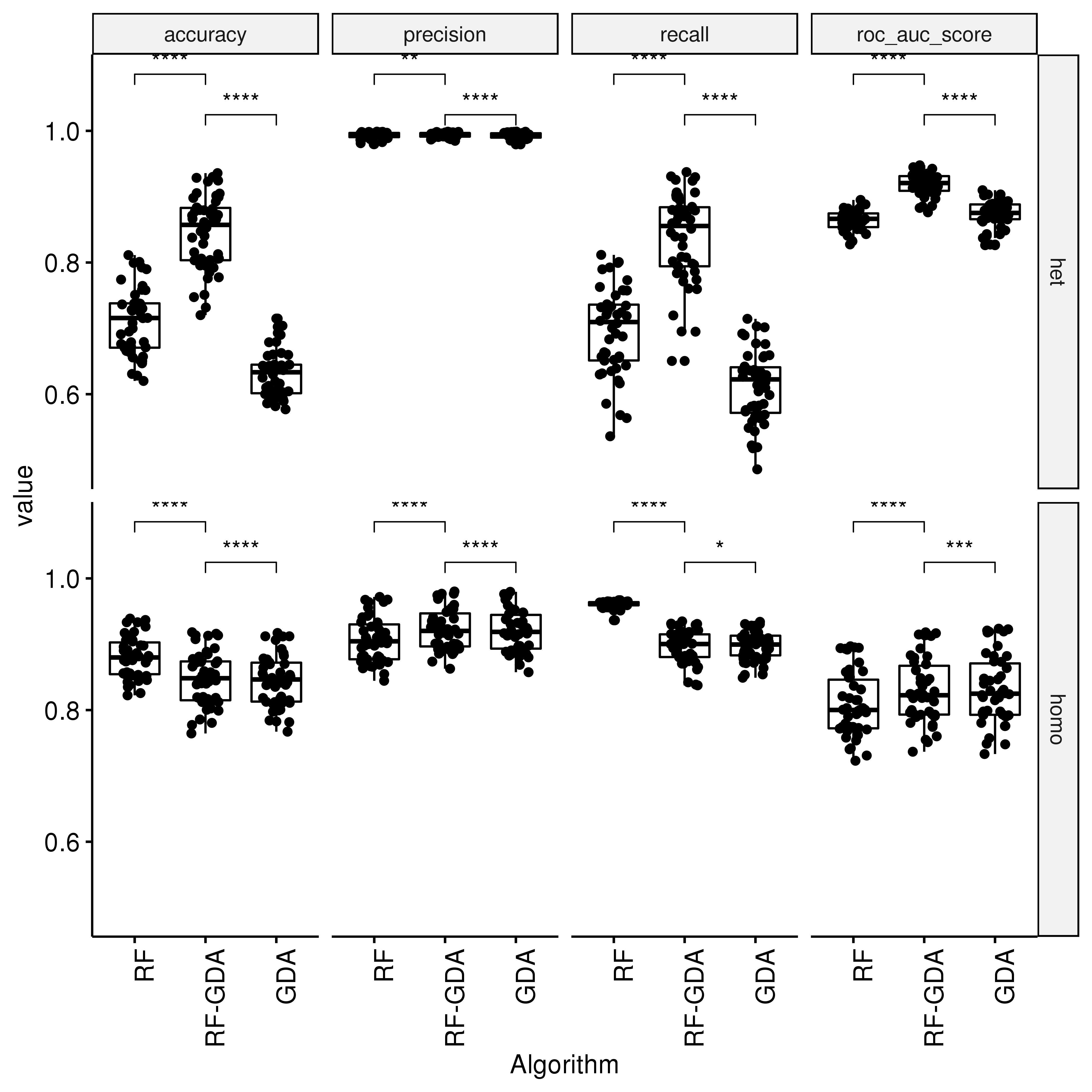

**Figure S5. Comparison of performance of the single layers (RF, GDA) vs. a combined two layered architecture (RF-GDA) on 46 cells from GM12878.** Each dot represents one cell. The pairwise statistics for the 46 single cells was performed using paired t-test.

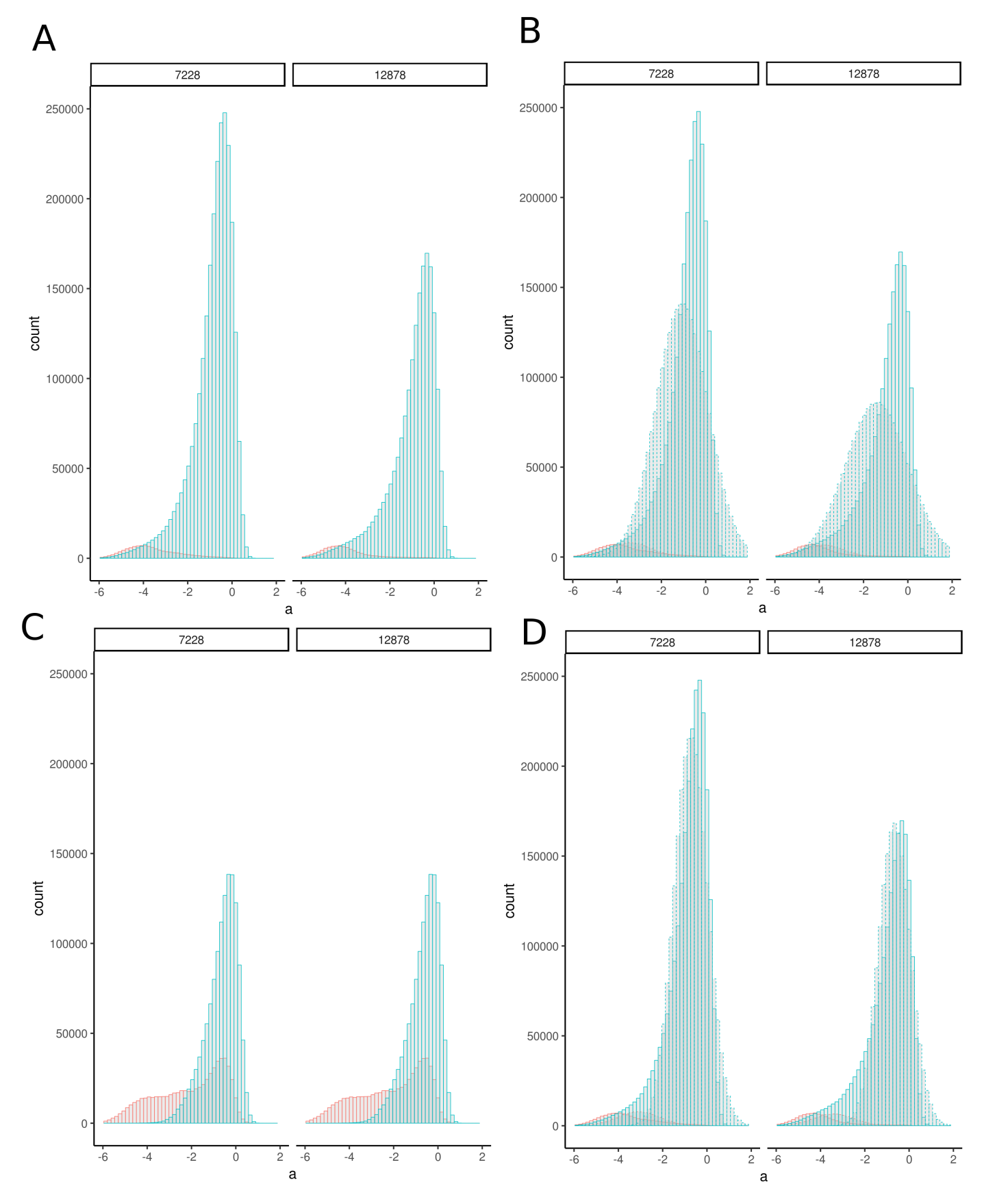

**Figure S6. Distribution of the a-feature (log average of intensities) from the heterozygous clusters.** Negative class (predicted or ground truth) is labeled in red, positive class in blue. (A) Empirical distribution from the ground truth data. (B) Empirical distribution (solid line) and distribution generated from the GDA model fitted on the ground truth (dashed line). (C) Empirical distribution of the data labeled by RF. (D) Empirical distribution from the ground truth (solid line) and distribution of the data generated from the GDA model trained on the RF prediction (dashed line).

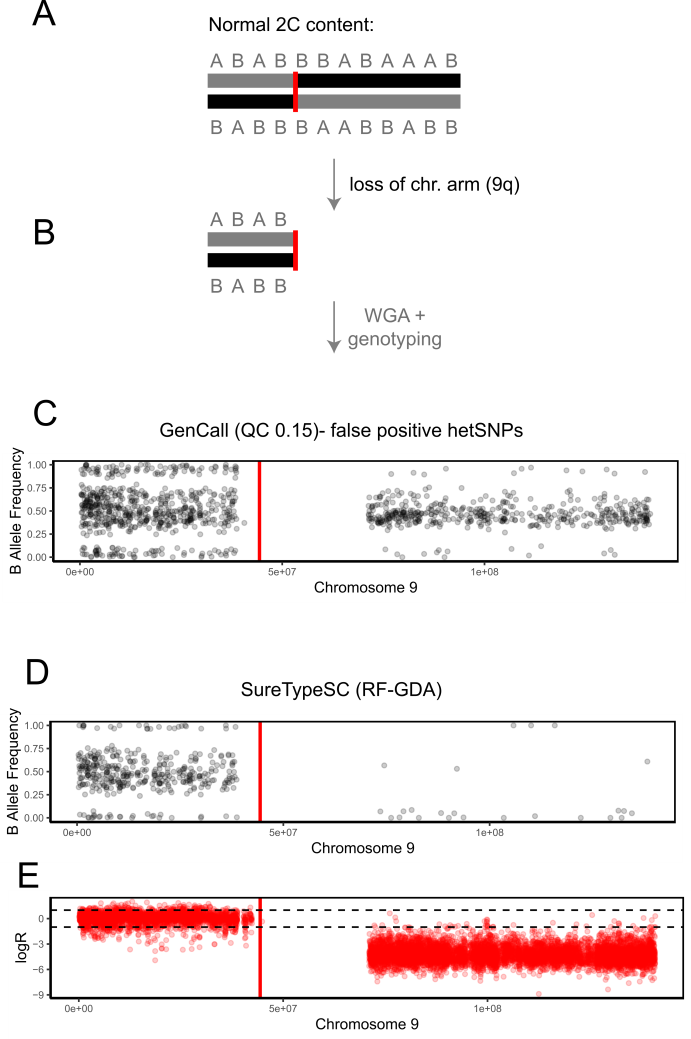

**Figure S7. Segmental aneuploidy in a human oocytes.** (A) The MII oocyte contains a haploid genome, with a is 2C content, but contains a partial deletion (the entire q arm). (B) After WGA, the genotyping algorithm should reject the signal noise from SNPs on the q arm. However, due to the normalization of SNP values, the B allele frequency of genotyped SNPs passed the quality control of GenCall (C). B allele frequency is inferred from the three canonical clusters from GenCall after an affine transformation causing the false positives of an entire chromosome arm. In contrast, the B allele frequency plot of SureTypeSC (D) correctly rejects genotypes from the deleted chromosome arms calling only a few SNPs. Figure (E) shows the logarithmic ratio of observed vs. expected signal intensities along the chromosome. It confirms that the area of the chromosomal loss has lower signal (negative logR).

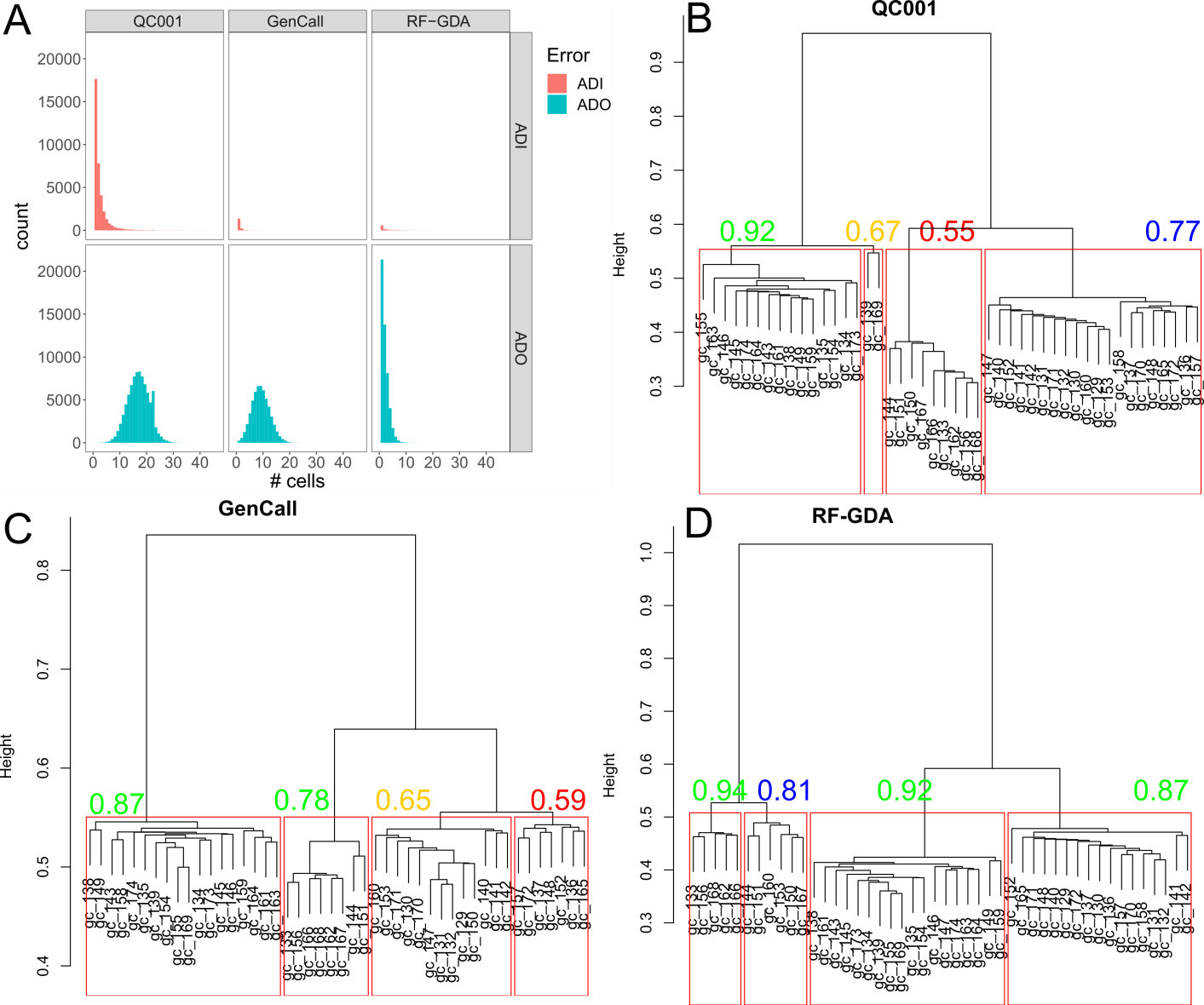

**Figure S8. Detected errors and potential variants in the 46 cells of GM12878.** (A) Histogram of detected ADI and ADO in raw data (left panel), data filtered with GenCall at high precision (GenCall score 0.87, middle panel) and data filtered with SureTypeSC (right panel). Hierarchical clustering on raw data (B), on data from GenCall (C) and from RF-GDA (SureTypeSC, D). The histograms were generated in R using hclust and evaluated using the clusterboot function from the R package *fpc.* The red rectangles show the potential subpopulations and the numbers indicate the Jaccard bootstrap mean. Labels are coloured according to the following rules (obtained from Henning 2007 and manual of the *fpc* package): Green labels indicate highly stable clusters (>0.85), blue labels indicate stable clusters (>0.75); orange labels might indicate a pattern, however the membership of the cells to a particular cluster is doubtful; red labels (<0.6) indicate unstable clusters. RF-GDA is the only algorithm that gives four stable clusters.

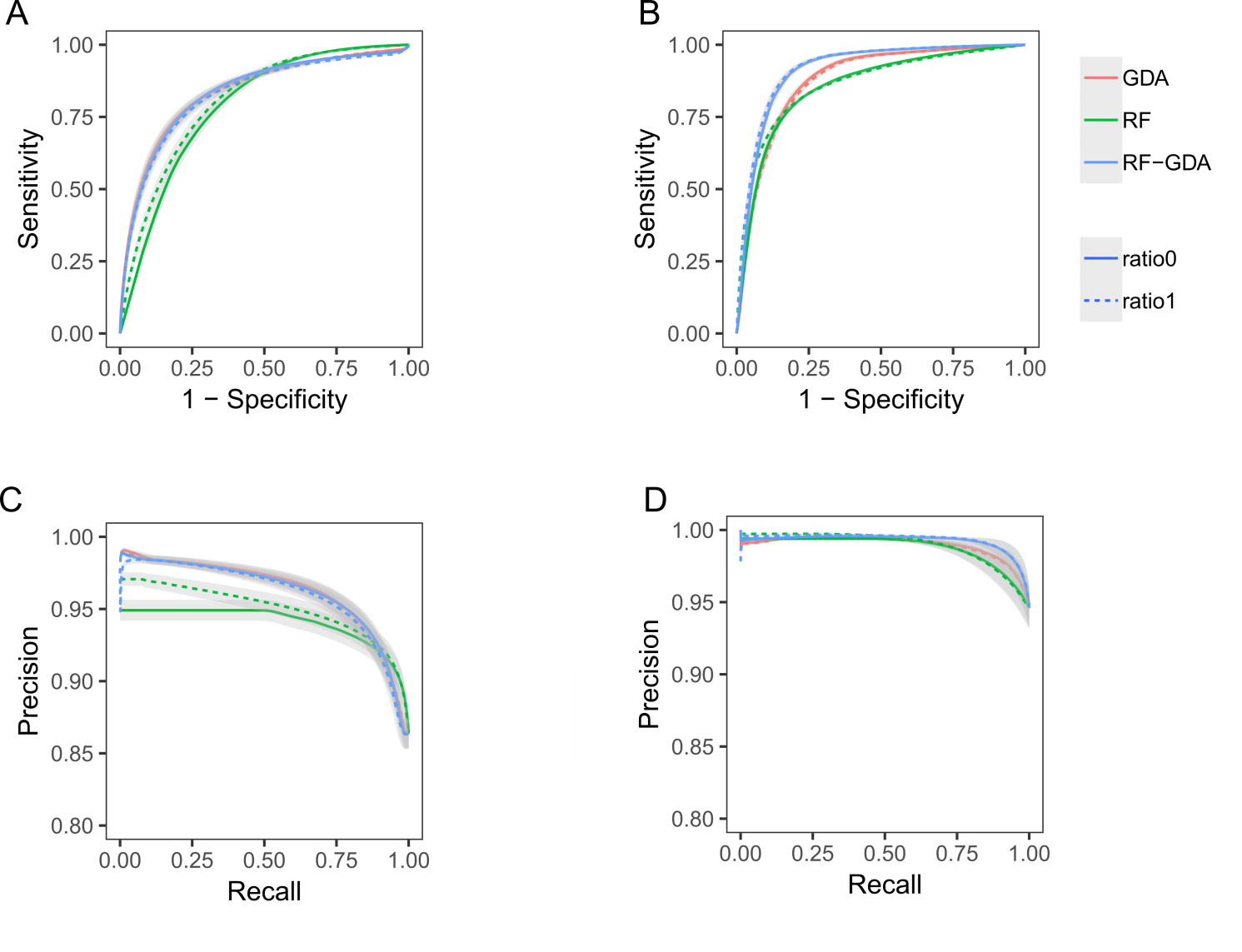

**Figure S9. Comparison of classifiers trained on balanced (dashed line) and unbalanced datasets (solid line).** ROC curve for the homozygous regions (A), ROC curve for the heterozygous regions (B). Precision recall curve for the homozygous regions (C) and for the heterozygous regions (D). Ratio0 refers to training data that reflect the biological ratio of correctly and incorrectly typed SNPs, which is highly unbalanced (ratio: 6.7). Ratio1 refers to training data where both classes are equally represented by downsampling the majority class (ratio: 1).
